# BARD1 links histone H2A Lysine-15 ubiquitination to initiation of BRCA1-dependent homologous recombination

**DOI:** 10.1101/2020.06.01.127951

**Authors:** Jordan R. Becker, Clara Bonnet, Gillian Clifford, Anja Groth, Marcus D. Wilson, J. Ross Chapman

## Abstract

Protein ubiquitination at sites of DNA double-strand breaks (DSBs) by RNF168 recruits BRCA1 and 53BP1, mediators of the homologous recombination (HR) and non-homologous end joining (NHEJ) DSB repair pathways, respectively. While NHEJ relies on 53BP1 binding to ubiquitinated Lysine 15 on H2A-type histones (H2AK15ub), an RNF168-dependent modification, the mechanism linking RNF168 to BRCA1 recruitment during HR has remained unclear. Here, we identify a tandem BRCT domain ubiquitin-dependent recruitment motif (BUDR) in BARD1 – BRCA1’s obligate partner protein – that binds H2AK15ub directly, thereby recruiting BRCA1 to DSBs. BARD1 BUDR mutations compromise HR, and render cells hypersensitive to PARP inhibition and cisplatin treatment. We find that BARD1-nucleosome interactions require BUDR binding to H2AK15ub and ankyrin repeat domain-mediated binding of the histone H4 tail, specifically when unmethylated on Lysine-20 (H4K20me0), a state limited to post replicative chromatin. Finally, we demonstrate that by integrating DNA damage-dependent H2AK15ub and DNA replication-dependent H4K20me0 signals at sites of DNA damage, BARD1 coordinates BRCA1-dependent HR with 53BP1 pathway antagonization, establishing a simple paradigm for the governance of DSB repair pathway choice.

---

The equilibrium between accurate DNA double-strand break (DSB) repair by homologous recombination (HR), and error-prone DSB repair by non-homologous end joining (NHEJ) is controlled by the BRCA1 and 53BP1 proteins and their interplay with two histone post-translational modification (PTM) states. DNA damage recognition by both proteins involves ubiquitination of H2A-type histones at DSB sites by the DNA damage responsive E3 ubiquitin ligase RNF168^1–4^. Chromatin engagement of 53BP1 and BRCA1 complexes also requires binding to histone H4 tails, through recognition of distinct lysine 20 methylation states that undergo DNA replication-dependent oscillations. Essential to its promotion of NHEJ, 53BP1 binds nucleosomes carrying histone H4 lysine 20 mono- and dimethylation (H4K20me1/2), histone PTMs highly abundant on old histones in pre- and post-replicative chromatin^5–7^. Conversely, BRCA1 complexes recognize H4 histones specifically when they are unmethylated at lysine 20 (H4K20me0), a state restricted to newly synthesized histones incorporated into chromatin during DNA replication^8^. H4K20me0 thereby recruits BRCA1 to post-replicative chromatin, where its promotion of HR is essential for genome stability and tumor suppression^4,8^.

Specialized histone binding domains in BARD1 (BRCA1-Associated RING Domain Protein 1) – BRCA1’s obligate interaction partner – and 53BP1, mediate histone H4 interactions. We recently showed the Ankyrin Repeat Domain (ARD) in BARD1 binds multiple residues in the H4 tail, and specifically Lysine-20 in its unmethylated state^8^, while the 53BP1 tandem-tudor domain (TTD) mediates the converse methylation-dependent interaction with H4K20me1/2^5^. To achieve specificity for chromatin proximal to DSBs, 53BP1 couples the binding of widespread H4K20me1/2 with recognition of the RNF168-dependent H2AK15ub PTM in DSB-proximal chromatin, using its ubiquitin-dependent recruitment motif (UDR), a TTD-proximal sequence that binds H2AK15ub and features of the nucleosome surface^9,10^. The equivalent dependence of BRCA1 recruitment on RNF168 activity^1,2^ similarly implicates H2AK15ub recognition, however the mechanism linking BRCA1 complexes to this modification during HR have remained unknown.

Given that 53BP1-nucleosome interactions involve simultaneous binding to H4K20me1 /2 and H2AK15ub, we considered whether the BRCA1-BARD1 complex might also possess sequences that bind H2AK15ub, and couple this to H4K20me0 recognition by the BARD1 ARD. BARD1 comprises an N-terminal RING, a central ARD, and a tandem BRCA1 C-terminal (BRCT) repeat domain at its C-terminus (Fig. 1a). Common to DNA damage responsive proteins, tandem BRCTs typically bind phosphoserine-containing peptide ligands in partner proteins^11–14^. Putative phosphopeptide-binding residues are conserved in the BARD1 BRCTs, yet reportedly bind to Poly-ADP-ribose (PAR) chains induced at sites of DNA damage^15^. Despite this, we recently showed a PAR-binding defective point mutant of BARD1 (BARD1^K619A^) was fully proficient in repairing olaparib-induced DNA lesions^8^. In agreement, mice homozygous for equivalent BARD1 BRCT mutations were not tumor prone and displayed a cellular proficiency for HR, dismissing a role for BRCT-dependent interactions with PAR or phospho-proteins in tumor suppression^16^. Nevertheless, when auxin-treated *BARD1^AID/AID^* HCT-116 cells – a cell-line engineered to encode biallelic auxin-dependent degron tags in the BARD1 C-terminus^8,17^ – were reconstituted with a BRCT domain-deleted BARD1 transgene (BARD1^ΔBRCT^), their hypersensitivity to olaparib (Fig. 1b) implicated their importance in HR^18^. Importantly, BARD1^ΔBRCT^ protein was expressed at endogenous levels and stabilized BRCA1 (Fig. 1c), prompting us to consider a specific and undescribed function for the BARD1 BRCTs in HR.

**Fig. 1.**
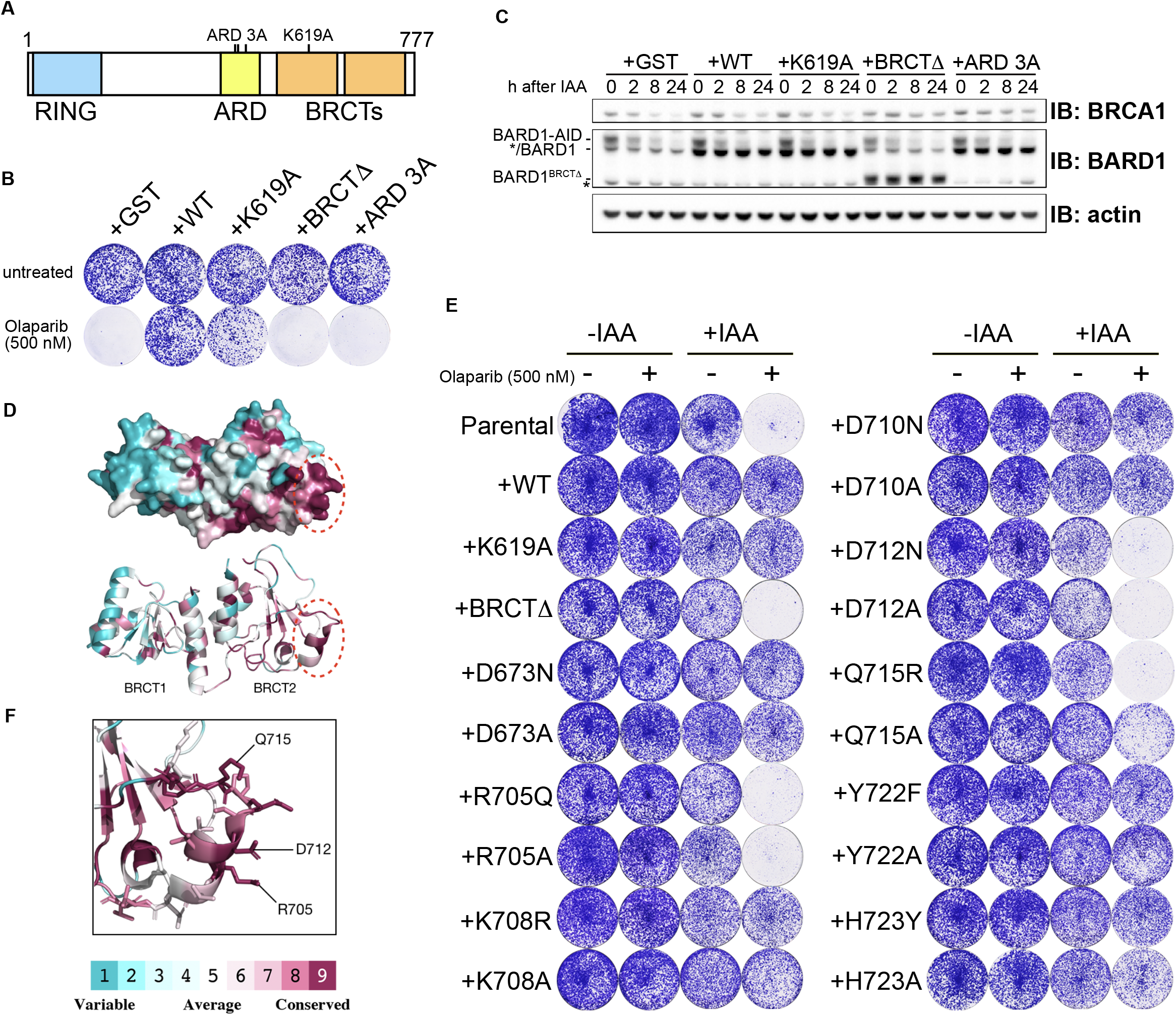
Residues in the inter-β2’-β3’ loop of BARD1 BRCT2 are essential for HR. **(A)** *BARD1* domain map with ARD 3A (N470A E467A D500A) and K619A mutations indicated. **(B)** Survival of indicated *BARD1^AID/AID^* cells grown in the presence of olaparib. Cell lines were grown in the presence of doxycycline (2 μg/ml) for 24 h before auxin addition (1 mM IAA). Olaparib (500 nM) was added 1 h after IAA. Cells were stained with crystal violet 10 days after the addition of olaparib. Representative data, *n=3* biological experiments. **(C)** Immunoblots of whole cell lysates harvested at the indicated timepoints after IAA addition. Expression of the auxin-degron-targeting SCF-complex E3 ligase OsTIR1 was induced using doxycycline (2 μg/ml), 24 h prior to the depletion of endogenous BARD-AID protein with IAA (1 mM). Representative of two biological repeats. **(D)** Space-filling (*top*) and ribbon (*bottom*) models of the BARD1 tandem BRCT crystal structure (PDB ID: 2NTE) pseudo-colored to indicate amino acid conservation. Red dashed ovals indicate the inter-β2’-β3’ loop. **(E)** As in (B). Representative data, *n=3* biological experiments. **(F)** The inter-β2’-β3’ loop with amino acid side-chains represented.

To identify putative functional surfaces in the BARD1 BRCTs, we mapped sequence conservation onto a crystal structure of this domain^19^, and used this to prioritize highly-conserved solvent-exposed residues for mutagenesis (Fig. 1d). BARD1 transgenes bearing neutral or disruptive amino acid substitutions at 9 positions were then stably integrated into *BARD1^AID/AID^* cells, and assayed for olaparib sensitivity following auxin-induced depletion of endogenous BARD1 (Fig. 1e and Extended Data Fig. 1a). Interestingly, only mutations within a focused cluster of 3 conserved residues – Arg-705, Asp-712 and Gln-715 – conferred olaparib sensitivity (Fig. 1e). These all mapped to the loop formed between beta-sheets 2 and 3 of BARD1’s second BRCT (inter-β2’-β3’ loop, Fig. 1f), a protruding feature comprising three 310 helices previously noted to be unique among BRCTs^19^. The observation that all three mutant BARD1 proteins were stable (Extended Data Fig. 1a-b), yet potentiated olaparib hypersensitivity, indicated a direct role for the inter-β2’-β3’ loop in HR.

We noted that BRCT_2_ inter-β2’-β3’ loop mutants exhibited olaparib sensitivity profiles equivalent to ARD mutated BARD1^ARD 3A^ expressing cell lines (Fig. 2a and Extended Data Fig. 2a), and considered that the BARD1 ARD and BRCTs might be functionally interconnected. Consistent with this notion, ARD 3A mutations did not synergize with the D712A inter-β2’-β3’ loop mutation in increasing cellular hyper-sensitivity to olaparib (Fig. 2b and Extended Data Fig. 2b) or cisplatin (Fig. 2c and Extended Data Fig. 2c). We thus used auxin-treated *BARD1^AID/AID^* cells complemented with either wild type, *BARD1^ARD 3A^, BARD1^D712A^*, or *BARD1^ARD3A/D712A^* double mutant BARD1 transgenes to assess whether cooperation between the ARD and tandem BRCT domains in BARD1 was necessary for BRCA1-BARD1 recruitment to DSB sites. High content imaging of BRCA1 ionizing radiation induced foci (IRIF) was combined with immunofluorescence intensity-labelling of H4-K20me0, to quantify BRCA1 recruitment in H4K20me0-high cell populations where IR-induced BRCA1 recruitment is highest^8^. Control (GST) complemented BARD1 deficient cells exhibited profound BRCA1 recruitment defects that were suppressed upon wild type BARD1 complementation (Fig. 2d-e and Extended Data Fig. 2d-e). However, only very low frequencies of BRCA1 IRIF were observed in cells complemented with the BARD1^ARD 3A^, BARD1^D712A^, or BARD1^ARD 3A/D712A^ transgenes (Fig. 2d-e and Extended Data Fig. 2d-e), confirming a requirement for ARD-BRCT cooperation in the recruitment of BRCA1. We suspected that residual BRCA1 IRIF detected in ARD and BRCT mutant-complemented cells were dependent on the BRCA1-A complex: a protein complex composed of BRCC36, ABRAXAS, BRE, MERIT40, and RAP80, which recruits BRCA1-BARD1 to DNA damage sites via RAP80-mediated interactions with Lysine-63-linked polyubiquitin chains^20–24^, yet is dispensable for BRCA1-dependent HR^25^. Consistent with previous reports^25,26^, BRCA1 IRIF frequencies in wild type BARD1 complemented *BARD1^AID/AID^* cells were only modestly reduced by deletion of RAP80 (Fig. 2f-g). By contrast, BRCA1 IRIF were ablated in *RAP8C^−/−^ BARD1^AID/AID^* cells complemented with the BARD1^D712A^, BARD1^ARD 3A^, and BARD1^ARD 3A,D712A^ double mutants (Fig. 2f-g and Extended Data Fig. 2f-g). Thus, the BARD1 ARD and BRCT domains recruit BRCA1 to DSBs independently of RAP80 and BRCA1-A. In confirmation of an essential role for ARD-BRCT interplay in HR, neither control (GST), BARD1^D712A^, BARD1^ARD 3A^ and BARD1^ARD 3A,D712A^-complemented *BARD1^AID/AID^* cells supported the recruitment of RAD51 into IRIF, in contrast to wild type BARD1-complemented cells, in which RAD51 frequencies were fully restored (Fig. 2h-i).

**Fig. 2.**
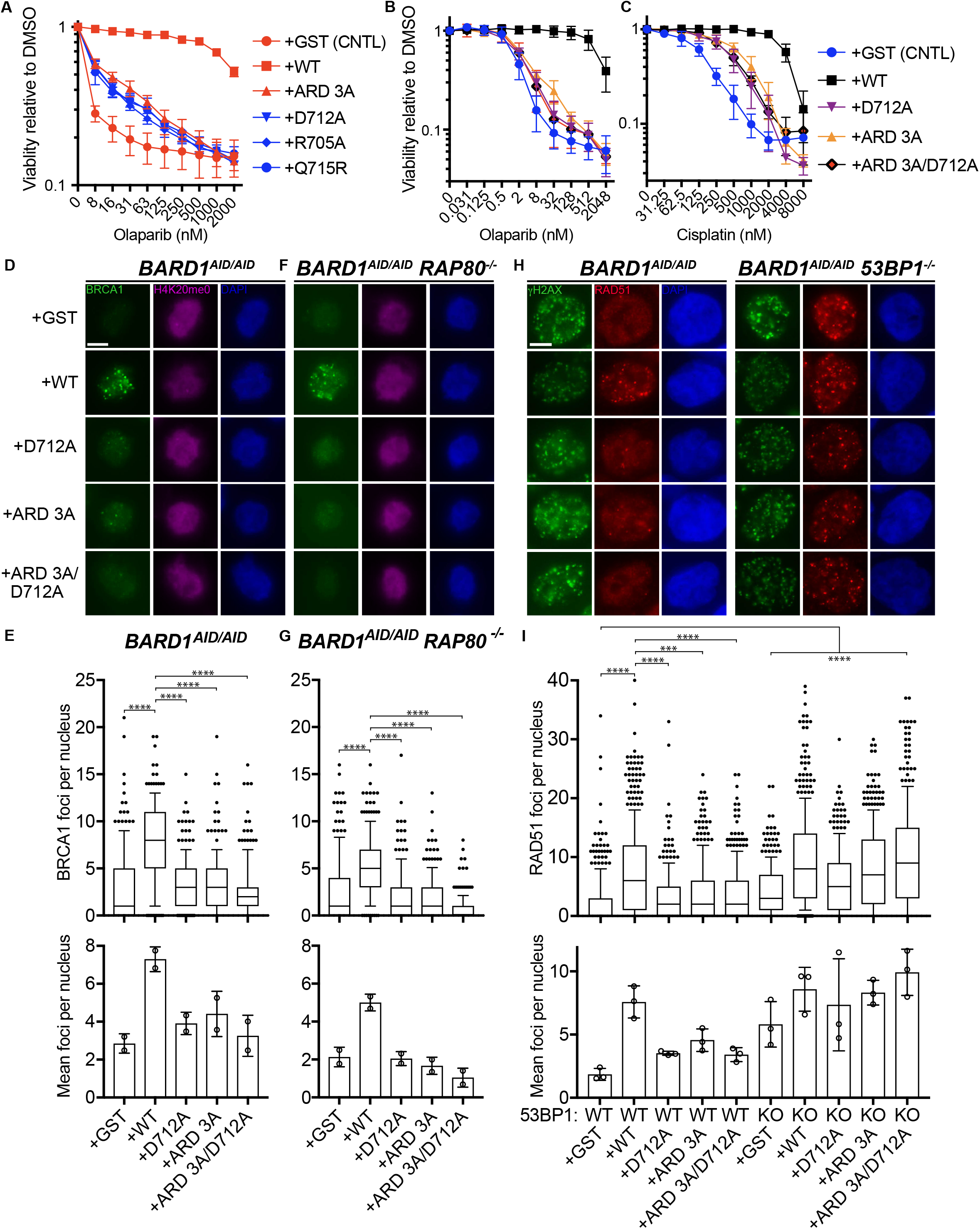
Ankyrin and BRCT repeat domains in BARD1 co-recruit BRCA1 during HR. **(A-C)** Survival of indicated *BARD1^AID/AID^* cell lines grown for 7 days in the presence of indicated doses of olaparib or cisplatin. Cell lines were seeded in doxycycline (2 μg/ml) for 24 h before IAA (1 mM) and olaparib or cisplatin addition. Resazurin cell viability assay, *n=3* biological experiments, mean ±s.d. **(D)** Immunofluorescent microscopy of BRCA1 IRIF in H4K20me0-positive *BARD1^AID/AID^* cell lines. Cultures were grown in the presence of doxycycline (2 μg/ml) for 24 h before IAA (1 mM) addition, irradiated 2 h later, and fixed with PFA 2 h following irradiation. Scale bar indicates 5 μm. Representative of *n=2* biological experiments. **(E)** *Top:* quantification of BRCA1 IRIF from (D). Boxes indicate the 25^th^-75^th^ percentiles with the median denoted and whiskers indicate the 10^th^-90^th^ percentiles. BRCA1 foci measurements are made for nuclei in the top quartile of H4K20me0 integrated staining intensity (>171 nuclei per condition). Integrated intensity and foci quantifications were made using CellProfiler. Significance was determined by Kruskal-Wallis H test with Dunn’s correction for multiple comparisons (****p < 0.0001). *Bottom:* Mean number of BRCA1 foci per cell from two independent experiments ±s.d. **(F and G)** Same as in (D-E) in *RAP80-^/^-* cells. >178 nuclei per condition. **(H)** Immunofluorescent microscopy of RAD51 IRIF in *BARD1^AID/AID^* cells expressing the indicated transgenes. Cultures were grown in the presence of doxycycline (2 μg/ml) for 24 h before IAA (1 mM) addition, irradiated 2 h later, and fixed with PFA 2 h following irradiation. Scale bar indicates 5 μm. Representative of *n=3* biological experiments. **(I)** *Top:* Quantification of Rad51 foci per cell from (H). >255 nuclei per condition. Significance was determined by Kruskal-Wallis H test with Dunn’s correction for multiple comparisons (****p ≤ 0.0001, ***p = 0.0003). Representative of *n=3* biological experiments. *Bottom:* Mean number of RAD51 IRIF from three independent biological experiments ±s.d.

Interdependence between the BARD1 ARD and BRCTs suggested their cooperation in chromatin binding at DSB sites. We therefore speculated the BARD1 C-terminal domain architecture might couple H4K20me0 and H2AK15ub binding in a manner analogous to the TTD-UDR domains of 53BP1. If the BARD1 BRCT repeats interacted with H2AK15ub, we reasoned that they might rescue the recruitment of a UDR mutated fragment of 53BP1 that encoded its minimal IRIF forming region (amino acids 1210-1711)^27^. We tested this hypothesis by expressing chimeric proteins in which wild-type or D712A mutant versions of the BARD1 BRCT repeats were fused C-terminal to wild-type or UDR-mutated fragments of 53BP1^aa1220-1711^, and examined their ability to form IRIF in *53BP1^−/−^ BARD1^AID/AID^* cells (Extended Data Fig. 3a). As expected, the 53BP1^aa1220-1711^:BARD^BRCT1-2^ fusion proteins readily formed IRIF that were completely ablated when recruitment-neutralizing L1619A^9,27^ and D712A mutations were introduced in the 53BP1 UDR and BARD1 BRCT repeats, respectively (Fig 3a). However, fusion of wild type BARD1 BRCTs to the UDR-mutant 53BP1 fragment rescued its recruitment into IRIF, indicating the BARD1 BRCT repeats can recruit BRCA1-BARD1 to the H2AK15ub PTM that demarks nucleosomes at DSB sites.

**Fig. 3.**
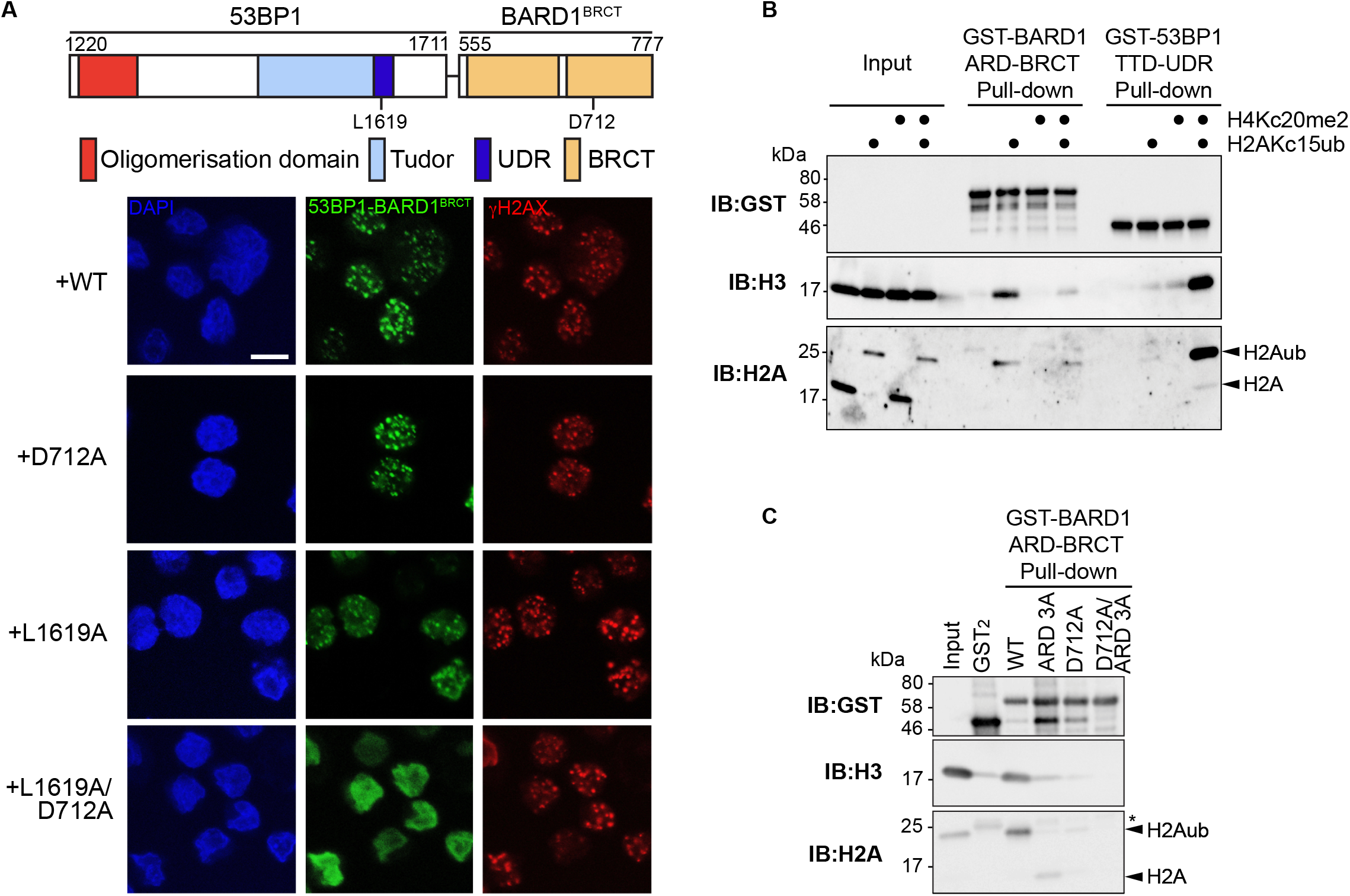
The inter-β2’-β3’ loop in BARD1 BRCT2 is a ubiquitin-dependent recruitment motif. **(A)** *Top:* Model depicting the 53BP1-BARD1 fusion protein. The fusion is a chimera composed of the 53BP1 minimal focus forming region (a.a. 1220-1711) and BARD1 BRCTs (a.a. 555-777). Expressed form includes an N-terminal 2xHA-FLAG epitope tag. *Bottom:* Confocal immunofluorescent microscopy of 53BP1-BARD1 chimeric fusion proteins in irradiated *BARD1^AID/AID^ 53BP1^−/−^* cells. Cultures were grown in the presence of doxycycline (2 μg/ml) for 24 h before IAA (1 mM) addition and irradiated (5 Gy) 2 h later. Cells were fixed 2 h following irradiation. Scale bar indicates 10 μm. Representative of *n=2* biological experiments. **(B)** Immunoblots from pull-down assay using GST tagged BARD1 (425-777) and 53BP1 (1484-1631) fragments immobilised on glutathione affinity beads, and incubated with recombinant nucleosome variants. Nucleosomes were either modified with dimethyl-lysine analogs at H4 position 20 and/or chemically ubiquitinated at H2A position 15. **(C)** Immunoblots from pulldown assay using tandem GST, GST tagged BARD1 (425-777) wild type (WT) and indicated point mutants. Pull-down was performed on immobilized GST tagged proteins incubated with recombinant nucleosomes ubiquitinated at position 15 on H2A. ARD 3A, comprises mutations (N470A E467A D500A) in the Ankyrin repeat domain while D712A is in tandem BRCT domain.

To directly test whether the BARD1 BRCT repeats, akin to the 53BP1 UDR, bind H2AK15ub labeled nucleosomes, GST fusion protein fragments encoding the BARD1^ARD-BRCT^ (a.a 425-777), or the 53BP1^TTD-UDR^ (a.a. 1484-1631) were recombinantly expressed and purified (Extended Data Fig. 3b). GST pull-downs were then performed following incubation with fully-defined reconstituted nucleosomes that were either unmodified, chemically methylated at H4K20 (H4Kc20me2), chemically ubiquitinated at H2AK15 (H2AKc15ub), or both (Extended Data Fig. 5c-d). As expected^9,10^, the presence of both histone PTMs stimulated nucleosome binding to 53BP1^TTD-UDR^ (Fig. 3b). In contrast, BARD1^ARD-BRCT^ binding to nucleosomes depended on the presence of H2AKc15ub alone, as interactions were inhibited when the H4Kc20me2 modification was also present (Fig. 3b). This confirmed that simultaneous binding of the BARD1 ARD-BRCT domains to H4K20me0 and H2AK15ub promotes robust BARD1-nucleosome interactions. Lastly, interactions between H2AKc15ub modified nucleosomes and GST-BARD1^ARD-BRCT^ fragments, were sensitive to the ARD 3A and D712A mutations alone, and in combination (Fig. 3c and Extended Data Fig. 3e). Thus, recruitment-defective mutations in either the ARD or BRCT domains block BARD1 binding to H2AK15ub-modified nucleosomes. We thus identify a critical and unique role for the BARD1 BRCT repeats in mediating ubiquitin-dependent histone interactions, prompting us to rename its inter-β2’-β3’ loop the BRCT ubiquitin-dependent recruitment motif (BUDR).

HR deficiency driven tumor predisposition in mice homozygous for hypomorphic *Brca1-*mutations is largely attributed to the 53BP1 pathway^28,29^. However, the phenotypic suppression conferred by 53BP1 deletion in mice homozygous for severely disruptive or nullizygous *Brca1* mutations is less pronounced, with double knockout cells retaining HR-defects and residual sensitivity to PARPi^30,31^. We therefore sought to determine whether HR defects accompanying loss of BARD1, or BARD1-dependent nucleosomal interactions could be entirely attributed to loss of 53BP1 pathway inhibition. In agreement, RAD51 IRIF were diminished in auxin-treated *BARD1^AID/AID^* cells, yet equally restored in *53BP1^−/−^ BARD1^AID/AID^* derivatives complemented with BARD1^ARD 3A^ or the BARD1^D712A^ BUDR mutant (Fig. 2h-i). Despite this, *53BP1^−/−^ BARD1^AID/AID^* cells retained significant sensitivity to both olaparib and cisplatin treatments (Fig. 4a-d), consistent with an incomplete HR restoration. However, re-expression of not only wild type BARD1, but also BARD1^D712A^ and BARD1^ARD 3A^ mutant proteins, fully restored cisplatin and olaparib resistance in *53BP1^−/−^ BARD1^AID/AID^* cells (Fig. 4a-d and Extended Data Fig. 4a-f). These findings demonstrate that 53BP1 pathway inhibition is the prime function for BARD1’s interaction with H4K20me0/H2AK15ub marked nucleosomes, a function consistent with BRCA1-dependent exclusion of 53BP1 from chromatin at DSB sites^8,32^. Our results also suggest that BRCA1-BARD1 complexes exert important functions that do not rely on BARD1 binding to chromatin, and whose loss cannot be alleviated by 53BP1 pathway loss. This may partly explain the incomplete phenotypic suppression by 53BP1-deletion in mice bearing severe *Brca1* loss-of-function alleles^30,31^.

**Fig. 4.**
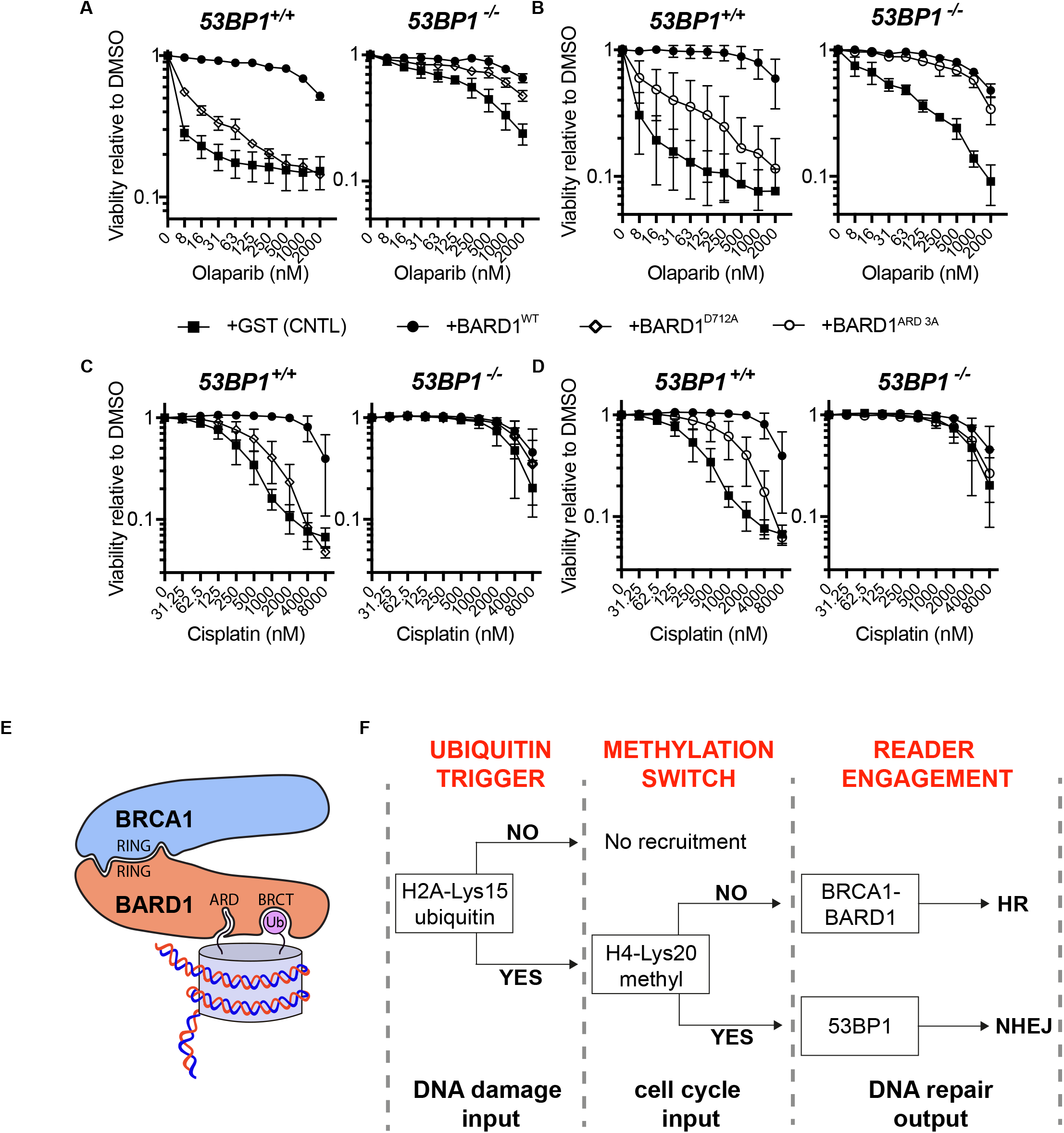
Two binary histone modifications and their readers control the repair fate of DSBs. **(A-D)** Survival of indicated *BARD1^AID/AID^* cell lines grown for 7 days in the presence of indicated doses of olaparib or cisplatin. Cultures were seeded in doxycycline (2 μg/ml). IAA (1 mM) was added after 24 h and olaparib or cisplatin was added 1 h after IAA. Survival was measured after 7 days by resazurin cell viability assay (*n=3* biological experiments) mean ±s.d. **(E)** Model of BRCA1-BARD1-nucleosome interactions. **(F)** Logic gate depicting how combinatorial H2AK15 and H4K20 PTM states govern DSB repair pathway choice.

Altogether, our results answer the long-standing question of how the DNA damage-associated H2AK15ub histone modification promotes BRCA1-BARD1 complex recruitment, to coordinate HR promotion with inhibition of 53BP1-dependent NHEJ. In identifying the BARD1 BRCT repeats as a receptor for this PTM, we also reveal a conserved and simple principle governing the equilibrium between competing DSB repair pathways, in which two histone PTM states – one cell-cycle regulated (H4K20±me1/2), and one DNA-damage dependent (H2AK15±ub) – can specify bivalent interactions with the reader domains of distinct repair pathway mediator protein (Fig 4e-f). The shared affinity of BARD1 ARD-BUDR and 53BP1 TTD-UDR architectures for H2AK15ub-modified nucleosomes, yet inverse affinities for H4-K20 methylation, explains the respective preferences of these proteins for DSB-associated chromatin in post- and pre-replicated regions of the genome, and the establishment of DSB repair pathway choice.

## Materials and Methods

### Cell lines and culture conditions

*BARD1^AID/AID^* cells lines were generated by biallelic knock-in of auxin-inducible degron tags at the C-terminus of the endogenous BARD1 loci in the adult male HCT116 colorectal carcinoma cells (parental cell line was a gift from I. Tomlinson, RRID: CVCL_0291) carrying doxycycline-inducible copies of OsTIR1 integrated the AAVS1 loci as previously described *(8).* All *BARD1^AID/AID^* and derivative cell lines were maintained in Dulbecco’s modified Eagle medium (DMEM)-high glucose (Sigma-Aldrich, D6546) supplemented with 10% FBS, Pen-Strep, and 2 mM L-glutamine. Cultures were maintained at 37°C with 5% CO_2_.

To generate lentivirus for stable transgene complementation, HEK 293T female embryonic kidney cells (RRID: CVCL_0063) were co-transfected with a lentiviral vector encoding the transgene of interest, pHDM-tat1b, pHDM-G, pRC/CMV-rev1b, and pHDM-Hgpm2 using 1.29 μg polyethylenimine per μg of DNA in Opti-MEM (Thermo Fisher, 31985062). Viral supernatants were harvested at 48 h and 72 h after transfection, syringe filtered (0.45-μm), and immediately used to transduce target cells populations in the presence of 4 μg/ml polybrene. Transduced populations were selected with antibiotic beginning 24 h after the last round of transduction until a non-transduced control population was completely dead. Stably transduced cell lines were maintained in the presence of selective antibiotic.

All knock-out cell lines were generated by CRISPR-Cas9. Gene-specific gRNAs were integrated into pSpCas9(BB)-2A-GFP (PX458) (Addgene #48138) and 2 μg of plasmid was electroporated into 10^6^ cells using a Lonza 4D-Nucleofector^TM^ according to the manufacturer’s protocol for HCT116 cells. GFP positive cells were sorted 24 h after electroporation using a Sony SH800 cell sorter with the brightest 5% being pooled for recovery in medium containing 50% FBS for 4 days. Sorted populations were then seeded at low density and individual clones were isolated after 10 days outgrowth. Individual clones were validated by western blot and sequencing.

### Survival experiments

To generate survival curves for *BARD1^AID/AID^* and derivative cell lines, 300 cells per well were seeded in the presence of doxycycline (2 μg/ml) in triplicate for each drug concentration in a 96-well plate. Each cell line was plated in duplicate for plus and minus IAA conditions. After 24 h, IAA (1 mM) or carrier (DMSO) was added. One hour following IAA addition, olaparib or cisplatin was added to the indicated final concentrations. Seven days after drug addition, the medium was replaced with phenol red-free DMEM (Thermo Fisher, 21063-029) supplemented with 10% FBS, Pen-Strep, 2 mM L-glutamine, and 10 μg/ml resazurin (Sigma-Aldrich, R7017). Plates were then returned to the incubator for 2-4 h or until the growth medium began to develop a pink color. Relative fluorescence was measured with a BMG LABTECH CLARIOstar plate reader. The mean of three technical repeats after background subtraction was taken as the value for a biological repeat and three biological repeats were performed for each experiment. All survival curves presented in this study represent the mean of three biological repeats ±s.d.

For survival experiments analyzed by crystal violet staining, 10^4^ cells were seeded per well of a 6-well plate in triplicate for each cell line in the presence of doxycycline (2 μg/ml). After 24 h, IAA (1 mM) or carrier (DMSO) was added. One hour following IAA addition, olaparib was added to the indicated final concentrations. Ten days after plating, the growth medium was removed and the cells were washed briefly with PBS before the addition of crystal violet stain (0.5% crystal violet in 25% methanol). Cells were stained for 5 minutes, washed with ddH2O and dried before scanning. Representative wells were selected for display.

### Immunofluorescence

For experiments analyzing BRCA1 foci, 10^6^ cells were passed through a 70 μm mesh cell strainer (Thermo Fisher, 22363548) and seeded in a single well of a 6-well plate in the presence of doxycycline (2 μg/ml). After 24 h, IAA was added to a final concentration of 1 mM. Cells were irradiated (5 Gy) 2 h after IAA addition, trypsinized 2 h after irradiation, and 10^5^ cells were plated on fibronectin-coated glass coverslips (13 mm) using a cytospin. Coverslips were immediately moved to ice cold cytoskeletal buffer (10mM PIPES pH 6.8, 300mM sucrose, 50mM NaCl, 3mM EDTA, 0.5% Triton X-100, Protease Inhibitor Cocktail [cOmplete™ EDTA-free; Roche, 27368400]) for 5 minutes before fixation in 2% PFA. *BARD1^AID/AID^ 53bp1^−/−^* cells stably transduced with 53BP1-BARD1 fusion protein were prepared identically as described for BRCA1 foci, but were immediately fixed in 2% PFA after cytospin. After fixation, these cells were permeabilized in PBS containing 0.2% Triton X-100.

We found RAD51 foci staining to be disrupted by cytospin plating. For RAD51 foci quantification, 2×10^5^ cells were passed through a 70 μm mesh cell strainer and seeded on 3 fibronectin-coated glass coverslips (13 mm) in a single well of a 6-well plate in the presence of doxycycline (2 μg/ml). After 24 h, IAA was added to a final concentration of 1 mM. Cells were irradiated (5 Gy) 2 h after IAA addition and fixed in 2% PFA 2 h after irradiation.

Staining of all fixed cells began with 15 min blocking (3% BSA, 0.1% Triton X-100 in PBS), followed by 1 h incubation with primary antibody in a humidity chamber. The following primary antibodies were used at the indicated concentrations: mouse anti-HA (1:200, HA.11 901501 Biolegend), mouse anti-BRCA1 D-9 (1:40, sc-6954 Santa Cruz), rabbit antiH4K20me0 (1:250, ab227804 Abcam), rabbit anti-RAD51 (1:1000, 70-001 BioAcademia), mouse anti-gH2AX (1:500, 05-636 Millipore), and rabbit anti-γH2AX (1:500, 2212-1 Epitomics). Following primary, coverslips were washed 3 times with PBS containing 0.1% Triton X-100 before incubation with secondary antibody for 1 h in a humidity chamber. Secondary antibodies used in this study were: goat anti-mouse Alexa Fluor 488 (1:500, A-11001 Invitrogen) and goat anti-rabbit Alexa Fluor 568 (1:500, A-11011 Invitrogen). Coverslips were then washed 3 more times with PBS containing 0.1% Triton X-100, once with PBS, and mounted on glass microscope slides using a drop of ProLong^®^ Gold antifade reagent with DAPI (Life Technologies, P36935).

Immunofluorescence images for quantification were acquired on a Leica DMi8 widefield microscope, while 53BP1-BARD1 fusion protein experiments were visualized on a Leica SP8-X SMD confocal microscope. CellProfiler (Broad Institute) was used for foci quantification. Images were visualized and saved in Fiji and assembled into figures in Adobe Illustrator.

### Protein extraction and western blotting

Cells were washed once with PBS and lysed by resuspension in ice cold benzonase cell lysis buffer (40 mM NaCl, 25 mM Tris pH 8.0, 0.05% SDS, 2 mM MgCl2, 10 U/ml benzonase, and cOmplete™ EDTA-free protease inhibitor cocktail [Roche, 27368400]). Extracts were then incubated on ice for 10 min before protein concentration was calculated by Bradford assay (Bio-Rad, 500-0006). Extracts were then mixed with Laemmli buffer and boiled at 95°C for 5 minutes before loading on SDS-PAGE gels.

Protein samples were fractionated on NuPAGE™ 4-12% 1.0 mm Bis-Tris polyacrylamide gels (Life Technologies, NP0322) before transferring to 0.45 μm nitrocellulose membranes (GE Healthcare, 10600003). After transfer, membranes with blocked with 5% milk in PBST for at least 30 min and then incubated overnight with primary antibody in PBST supplemented with 0.03% NaN_3_ and 3% BSA. Primary antibodies used for western blot in this study include: rabbit anti-53BP1 (Novus Biological, NB100-304, 1:2500), mouse anti-BRCA1 D-9 (1:400, sc-6954 Santa Cruz), rabbit anti-BARD1 (1:500, ab64164 Abcam), mouse anti-β-actin (1:2000, A1978 Sigma-Aldrich), and mouse anti-HA (1:2000, HA.11 901501 Biolegend). Following primary, membranes were incubated with either HRP-conjugated goat anti-mouse (1:20,000, Thermo Fisher, 62-6520) or HRP-conjugated goat anti-rabbit (1:20,000, Thermo Fisher, 65-6120) secondary antibodies. Membranes were developed with Clarity™ Western ECL Substrate (Bio-Rad, 170-5061) and imaged using a Gel Doc™ XR System (Bio-Rad).

For nucleosome pull-down assays, proteins were separated using 4-20% tris glycine gradient gels (BioRad) prior to transfer onto PVDF membranes. All blocking and antibody incubations were performed in Tris-buffered saline containing either 5% (w/v) BSA or 5% (w/v) skimmed milk powder. For Western blotting the following commercial primary antibodies were used: rabbit anti-H2A (Abcam, ab18255), rabbit anti-H3 (Abcam, ab1791), mouse anti-GST (Santa Cruz, sc-138). HRP-conjugated goat anti-rabbit IgG (Vector Laboratories, PI-1000) and HRP-conjugated horse anti-mouse IgG (Vector Laboratories, PI-2000) secondary antibodies were used with enhanced chemiluminescence solution (ECL supersignal, Thermo Scientific) was used for protein detection.

### Protein purification

GSTx2 and GST-BARD1 ANKD-BRCT variants were expressed using 200uM IPTG in BL-21 DE3 RIL E.coli overnight cultures grown at 16°C in 2YT broth. Cell pellets were resuspended in lysis buffer (25mM Tris pH 7.5, 300mM NaCl, 0.1% Triton (v/v), 10% glycerol (v/v), 5 mM β-mercaptoethanol, 1× Protease Inhibitor mix [284 ng/ml leupeptin, 1.37 μg/ml pepstatin A, 170 μg/ml phenylmethylsulfonyl fluoride and 330 μg/ml benzamindine], 1mM AEBSF and 5 μg/ml DNaseI). Cells were lysed by sonication and lysozyme treatment and spun at 39000g for 30 minutes. Clarified lysate was applied to a Glutathione Sepharose 4B (GE Healthcare). After extensive washing, bound protein was eluted using 30mM reduced glutathione and concentrated a 30K MWCO centrifugation device (Amicon). GSTx2 and GST-BARD1 ANKD-BRCT variants were further purified by size exclusion chromatography using a Superdex 200 Increase 10/300 in SEC buffer (20mM Tris pH 7.5, 150mM NaCl, 1mM DTT, 5 % glycerol) and the main mono-disperse protein containing peak was collected, concentrated, flash frozen in liquid nitrogen and stored at −80°C. GST-53BP1 Tudor-UDR was expressed and purified as described ^10^.

Protein concentrations were determined via absorbance at 280 nm using a Nanodrop 8000 (Thermo Scientific), followed by SDS-PAGE and InstantBlue (Expedeon) staining with comparison to known amounts of control proteins (Extended Data Fig. 3b and 3e).

Human histone proteins including site specific cysteine mutations were expressed in BL-21 DE3 RIL cells and purified from inclusion bodies, essentially as described^10,33^. 6-His-TEV-ub G76C was expressed in *E. coli* BL-21 DE3 CodonPlus cells, lysed in 1×Recom-500 buffer (25 mM Na-Phosphate buffer pH 7.4, 300 mM NaCl, 0.1 % [v/v] Triton, 10 % [v/v] glycerol, 4 mM β-mercapthoethanol, 1× Protease Inhibitor mix, 5 μg/ml DNaseI) and treated with lysozyme and sonication. Clarified cell lysate was loaded onto a HiTrap chelating column (GE Healthcare) pre-loaded with Ni^2+^ ions. After extensive washing, 6xHis-TEV-ub was eluted using a gradient of imidazole and peak-protein containing fractions were concentrated using a 3K MWCO centrifugation device (Amicon). 6xHis-TEV-ub was further purified on a S75 10/300 column in SEC buffer. Protein-containing fractions were dialyzed into water supplemented with 1 mM acetic acid prior to lyophilization.

### H4 Methyl lysine analog preparation

H4K20C was expressed and purified as described for other histones. Cysteine-engineered histone H4 K20C protein was alkylated essentially as described^34^. Briefly pure histone H4 was reduced with DTT prior to addition of a 50-fold molar excess of (2-chloroethyl) dimethylammonium chloride (Sigma-Aldrich). The reaction was allowed to proceed for four hours at room temperature before quenching with 5 mM β-mercaptoethanol. The H4 protein was separated and desalted using a PD-10 desalting column (GE Healthcare), preequilibrated in water supplemented with 1mM acetic acid and lyophilized. After incorporation of alkylation agents was assessed by 1D intact weight ESI mass spectrometry, roughly 85% was found to be modified. Lyophilized H4 was subject to a second round of alkylation as described above with the final reaction proceeding to near completion (>95%).

### H2A chemical ubiquitylation

Mutant human histone H2A engineered with a single cross-linkable cysteine (H2A K15C) was chemically ubiquitylated essentially as described^10,35^. Briefly, an alkylation reaction was assembled with H2A K15C (700uM), 6xHis-TEV-ubiquitin G76C (700uM) and 1,3-dibromoacetone (4.2mM, Santa Cruz) in 250 mM Tris-Cl pH 8.6, 8 M urea and 5 mM TCEP and allowed to react for 16 hours on ice. The reaction was quenched by the addition of 10 mM β-mercaptoethanol and pH adjusted to 7.5. Chemically ubiquitylated H2A (H2A Kc15ub) was purified using a HiTrap SP HP column (GE Healthcare) and 6xHis-TEV-H2AKc15ub containing fractions were pooled and enriched over a HiTrap chelating column (GE Healthcare) pre-loaded with Ni^2+^ ions. The 6xHis tag was removed by TEV cleavage and subsequent Ni^2+^ column subtraction. The resulting flow-through was dialysed against a 2mM β-mercaptoethanol/dH20 solution and lyophilized. H2AKc15ub was refolded and wrapped into nucleosomes as described below.

### Nucleosome reconstitution

Nucleosomes were reconstituted essentially as described^10,33^. Biotinylated 175bp Widom-601 DNA fragments for wrapping nucleosomes were generated by PCR based amplification, essentially as described^36^. For PCR amplification, 384 100ul reactions PCR reactions using Pfu polymerase and HPLC pure oligos (IDT) were pooled, filtered and purified using a ResourceQ column and salt gradient.

For octamer formation, 4 core histones were mixed at equimolar ratios in unfolding buffer (7M Guanidine HCl, 20mM Tris pH 7.5, 5mM DTT) prior to dialysis to promote refolding into 2M NaCl, 15mM Tris pH 7.5, 1mM EDTA, 5mM β-mercaptoethanol. Octamers were selected by gel filtration chromatography and assembled into nucleosomes via salt gradient dialysis. Soluble nucleosomes were partially precipitated with 9% polyethylene glycol (PEG) 6000 and resuspended in 10mM HEPEs pH7.5, 100mM NaCl, 1mM EDTA, 1mM DTT. Nucleosome formation and quality was checked by native gel electrophoresis and used within one month of wrapping (Extended Data Fig. 3d).

### Nucleosome Pull-down assays

Pull-down assays were performed essentially as described^10^. Briefly, 2.5 μg of GST-tagged 53BP1 or 8.5 ug of GST-BARD1/GST2 was immobilized on BSA-blocked Glutathione sepharose beads. Beads were separated and incubated with 2.2ug of nucleosome variant in pull-down buffer (50 mM Tris-HCL pH 7.5, 150mM NaCl, 0.02% NP40, 0.1 mg/ml BSA, 10 % glycerol, 1mM EDTA, 2mM β-mercaptoethanol) for 2 hours with rotation at 4°C. Pull-downs were washed three times in pull-down buffer and resuspended directly in 2× SDS loading buffer. All pull-down assays were repeated at least two times, with a single representative immunoblot displayed.

### Statistics

Prism 7 (Graphpad Software Inc.) was used for graphing and statistical analysis. Relevant statistical methods for individual experiments are detailed within figure legends.

## Supporting information

Compiled Extended Data Figures

## Acknowledgments

We thank all members of the Chapman lab for discussions, and Daniel Durocher for plasmid reagents. We acknowledge the BHF Centre of Research Excellence, Oxford (RE/13/1/30181), and the WHG Cellular Imaging core for equipment access and support. This work was funded by Cancer Research UK (CRUK) through the CRUK Oxford Centre (C5255/A18085 to JRB and JRC), and a CRUK Career Development Fellowship (C52690/A19270 to JRC) which also provided salary support to JRB. JRB’s salary is currently provided by a Ruth L. Kirschstein NRSA Individual Postdoctoral Fellowship (F32) (NIH/NCI – 1F32CA239339). CB was sponsored by an ERASMUS+ internship. MDW’s work is supported by Wellcome [210493] and the University of Edinburgh. The Wellcome Centre for Cell Biology is supported by core funding from the Wellcome Trust [203149]. The Wellcome Centre for Human Genetics is supported by Wellcome core award 090532/Z/09/Z. AG’s research is supported by the European Research Council (ERC CoG no. 724436), Independent Research Fund Denmark (7016-00042B), the Novo Nordisk Foundation (NNF14CC0001; NNF14OC0012839), and the Lundbeck Foundation (R198-2015-269).

## Author Contributions

JRB designed and analyzed the majority of experiments, and supervised experiments and analysis performed by CB. MDW and GC undertook and analyzed all *in vitro* nucleosome binding experiments. The project was initiated in collaboration with AG. JRC conceived and supervised the project and designed experiments. JRB and JRC co-wrote the manuscript, with input from authors.

## Data and materials availability

All data is available in the main text or extended data. Material requests should be addressed to JRC.

