## Supplementary material for "BARD1 links histone H2A Lysine-15 ubiquitination to initiation of BRCA1-dependent homologous recombination": Compiled Extended Data Figures

### EXTENDED DATA FIGURE 1

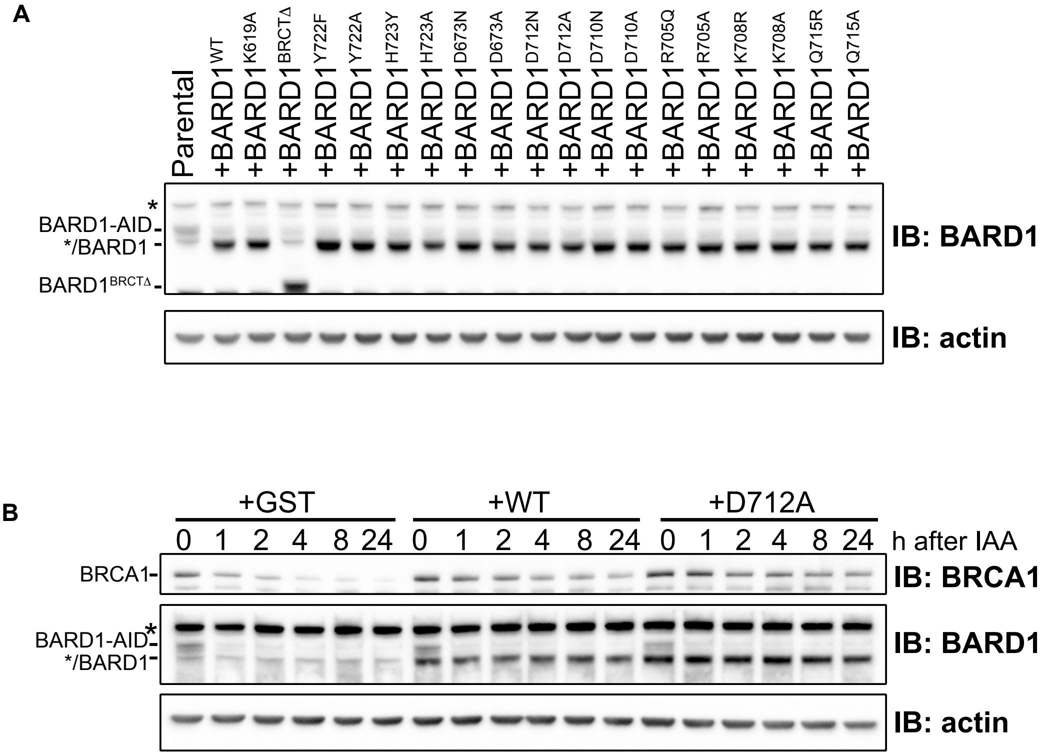


**Extended Data Fig. 1. BARD1 BRCT-mutated transgenes are stably expressed in *BARD1$^{AID/AID}$* HCT-116 cells (related to Figure 1). (A)** Immunoblot from whole cell lystes of BARD1 BRCT mutants screened for olaparib sensitivity. **(B)** Immunoblot of whole cell lysates from *BARD1$^{AID/AID}$* expressing the indicated transgenes. Cells were seeded in the presence of doxycycline (2 μg/ml) and IAA (1 mM) was added after 24 h. Lysates were harvested at the indicated timepoints after IAA addition. Representative of two biological repeats.

### EXTENDED DATA FIGURE 2

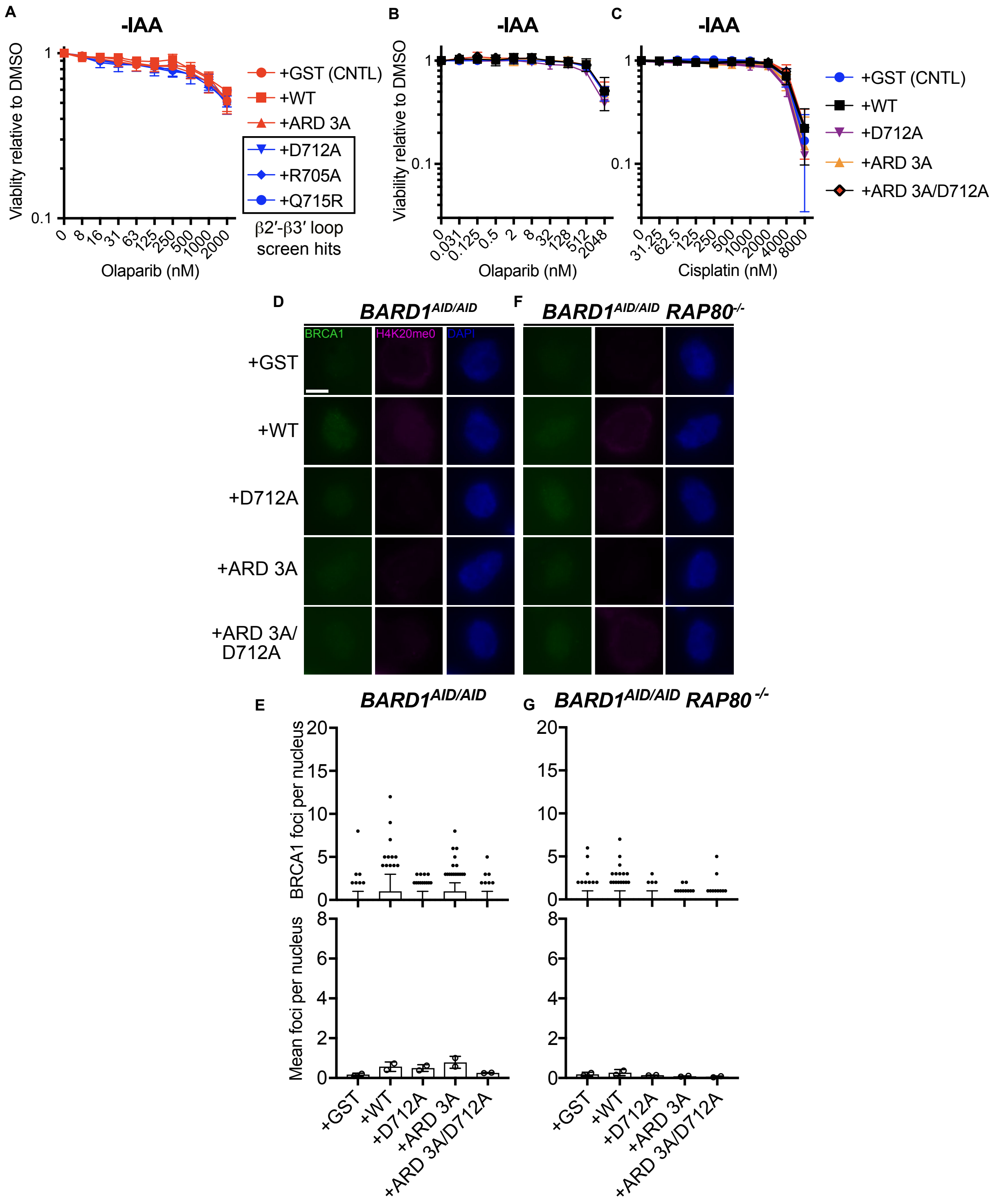

**Extended Data Fig. 2. Ankyrin and BRCT repeat domains in BARD1 cooperate in recruiting BRCA1 to post-replicative chromatin during HR. (Related to Figure 2). (A-C)** Survival of indicated *BARD1*$^{AID/AID}$ cell lines grown for 7 days without IAA in the presence of indicated doses of olaparib or cisplatin. Cell lines were seeded in doxycycline (2 μg/ml) for 24 h before olaparib or cisplatin addition. Resazurin cell viability assay, *n=3* biological experiments, mean ±s.d. **(D)** High-content immunofluorescent microscopy of BRCA1 IRIF in H4K20me0-positive *BARD1*$^{AID/AID}$ cells expressing the indicated transgenes. Cultures were grown in the presence of doxycycline (2 μg/ml) for 24 h before IAA (1 mM) addition, irradiated 2 h later, and fixed with PFA 2 h following irradiation. Scale bar indicates 5 μm. Representative of *n=2* biological experiments. **(E)** *Top*: quantification of BRCA1 IRIF from (D). Boxes indicate the 25$^{th}$-75$^{th}$ percentiles with the median denoted and whiskers indicate the 10$^{th}$-90$^{th}$ percentiles. BRCA1 foci measurements are made for nuclei in the bottom quartile of H4K20me0 integrated staining intensity (≥172 nuclei per condition). Integrated intensity and foci quantifications were carried out using CellProfiler. *Bottom:* Mean number of BRCA1 foci per cell from two independent experiments ±s.d. **(F and G)** Same as in (D and E) in *rap80*$^{-/-}$ cells. ≥179 nuclei per condition.

#### EXTENDED DATA FIGURE 3

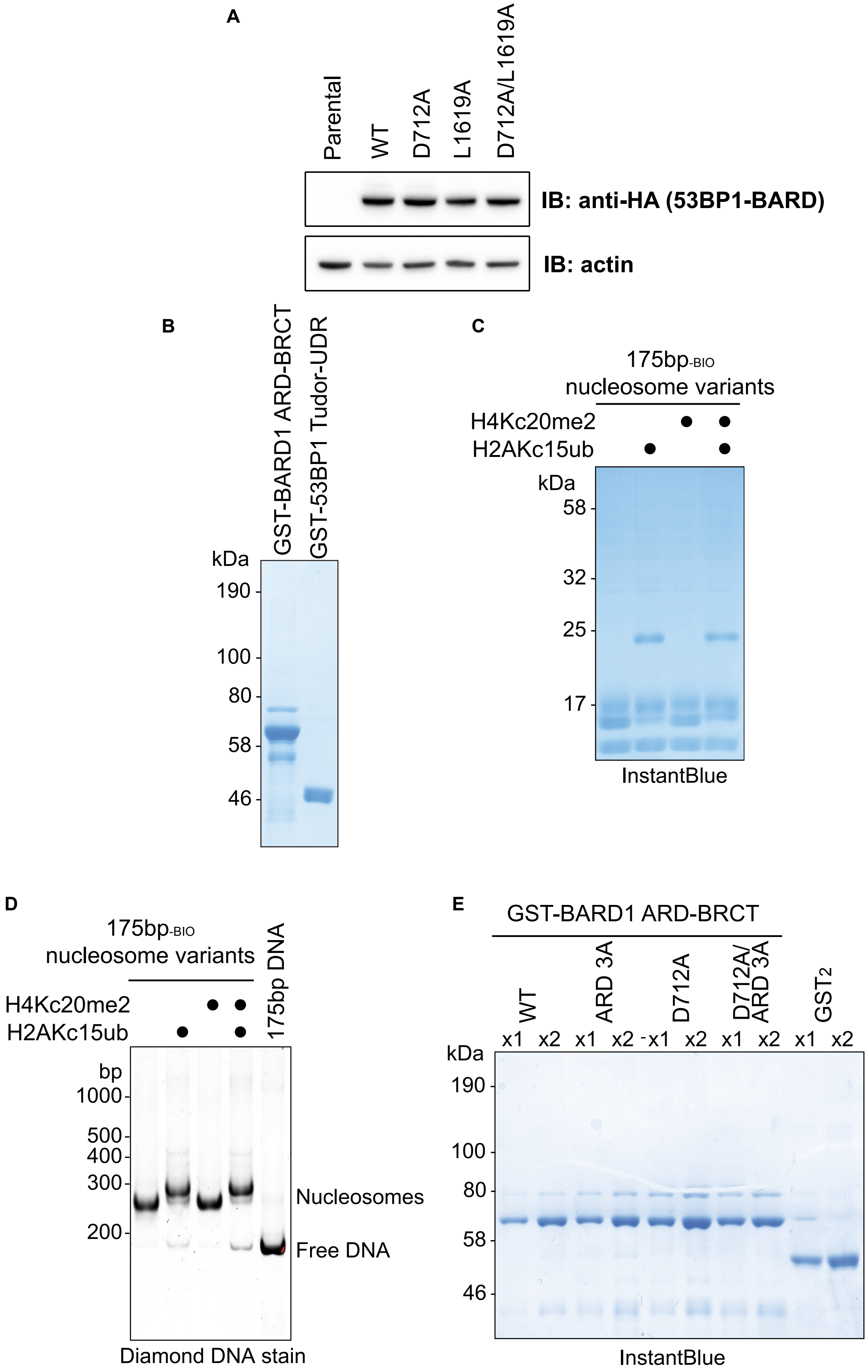

**Extended Data Fig. 3. Purification of BARD1 and 53BP1 fragments and assembly of modified nucleosomes. (A)** Western blot of HA-tagged 53BP1-BARD1 fusion proteins used in Fig. 5A stably expressed in *BARD1*$^{AID/AID}$ *53bp1*$^{-/-}$ cells. **(B)** SDS-PAGE gel, stained with InstantBlue protein stain of proteins used in Fig. 5B. **(C)** SDS-PAGE gel, stained with InstantBlue protein stain of nucleosomes used in this study. **(D)** Native gel electrophoresis of Widom 601 DNA in isolation and wrapped with nucleosomes used in this study. **(E)** SDS-PAGE gel, stained with InstantBlue protein stain of BARD1 variants used in Fig. 5C. Neighbouring lanes were loaded with two different concentrations.

### EXTENDED DATA FIGURE 4

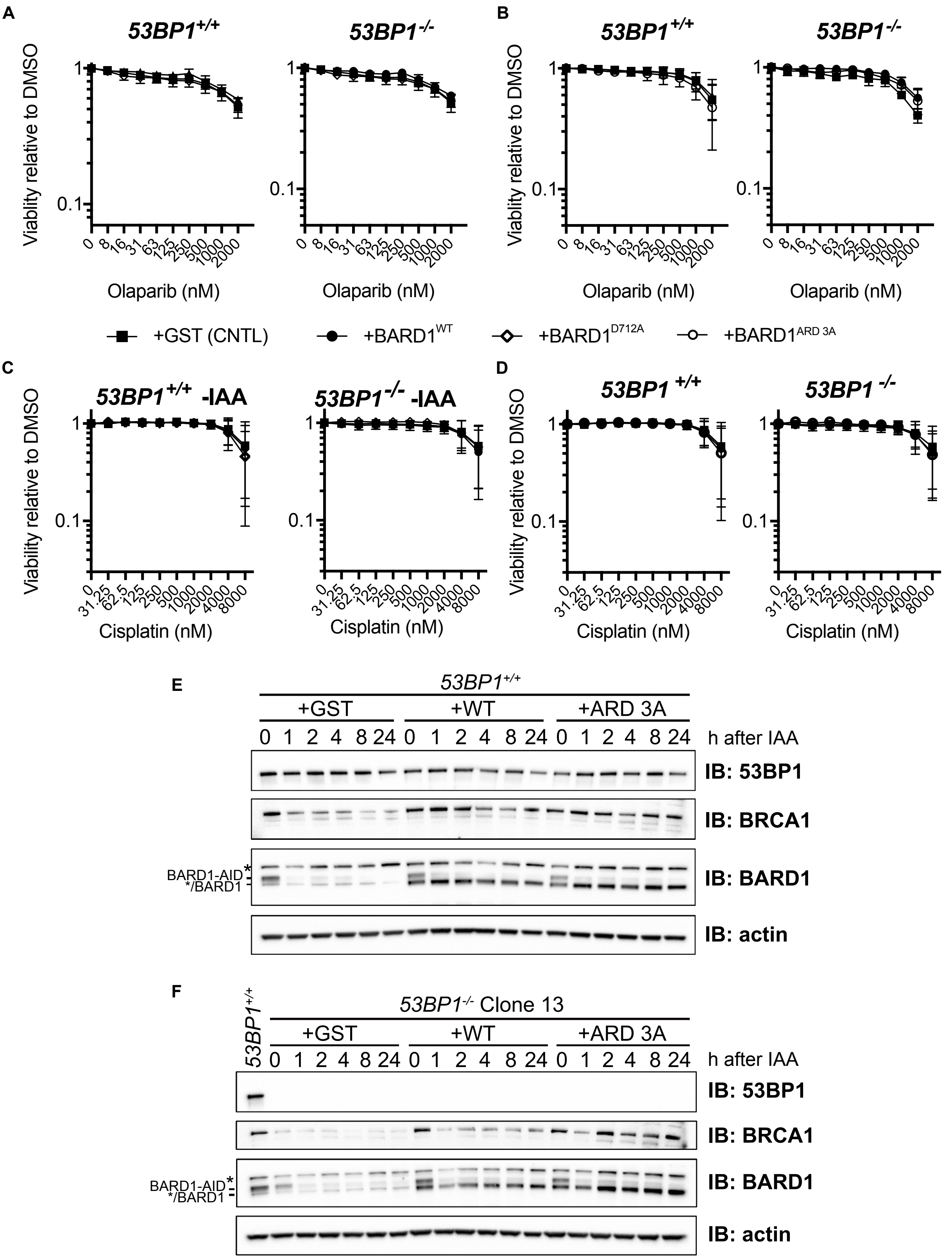

**Extended Data Fig. 4. The β2′-β3′ loop and ARD counteract toxic 53BP1-dependent NHEJ (related to Figure 4). (A-D)** Survival of indicated *BARD1*$^{AID/AID}$ cell lines grown without IAA for 7 days in the presence of indicated doses of olaparib or cisplatin. Cultures were seeded in doxycycline (2 μg/ml) and olaparib or cisplatin was added 24 h later. Survival was measured after 7 days by resazurin cell viability assay (*n=3* biological experiments) mean ±s.d. **(E and F)** Immunoblot of whole cell lysates from *BARD1*$^{AID/AID}$ (top) or *BARD1*$^{AID/AID}$ *53BP1*$^{-/-}$ (*bottom*) cells expressing the indicated transgenes. Cells were seeded in the presence of doxycycline (2 μg/ml) and IAA (1 mM) was added after 24 h. Lysates were harvested at the indicated timepoints after IAA addition. Representative of two biological repeats.
